# Pan-cancer deconvolution of cellular composition identifies molecular correlates of antitumour immunity and checkpoint blockade response

**DOI:** 10.1101/281592

**Authors:** Ankur Chakravarthy, Andrew Furness, Kroopa Joshi, Ehsan Ghorani, Kirsty Ford, Matthew J Ward, Emma V King, Matt Lechner, Teresa Marafioti, Sergio Quezada, Gareth J Thomas, Andrew Feber, Tim R Fenton

## Abstract

The nature and extent of immune cell infiltration into solid tumours are key determinants of therapeutic response. Here, using a novel DNA methylation-based approach to tumour cell fraction deconvolution, we report the integrated analysis of tumour composition and genomics across a wide spectrum of solid cancers. Initially studying head and neck squamous cell carcinoma, we identify two distinct tumour subgroups: ‘immune hot’ and ‘immune cold’, which display differing prognosis, mutation burden, cytokine signalling, cytolytic activity, and oncogenic driver events. We demonstrate the existence of such tumour subgroups pan-cancer, link clonal-neoantigen burden to hot tumours, and show that transcriptional signatures of hot tumours are selectively engaged in immunotherapy responders. We also find that treatment-naive hot tumours are markedly enriched for known immune-resistance genomic alterations and define a catalogue of novel and known mediators of active antitumour immunity, deriving biomarkers and potential targets for precision immunotherapy.

## Introduction

The tumour microenvironment plays key roles in shaping tumour evolution and in determining treatment responses; prominent intratumoural lymphocyte infiltration is a favourable prognostic marker in multiple tumour types^1–5^, while a high stromal content of extracellular matrix-producing cancer-associated fibroblasts (CAF), is associated with poor outcomes^6–8^. The recent clinical success of immunotherapy in subpopulations of patients with previously intractable malignancies has also highlighted the importance of understanding the tumour microenvironment in order to identify those patients who will derive most benefit from targeted therapies^9^. Although responses to immune checkpoint modulation (e.g. antibodies against PD-1 (programmed cell death protein 1), PD-L1 (programmed death-ligand 1 and CTLA-4 (cytotoxic T-lymphocyte-associated protein 4)) are seen across many solid tumours, the proportion of patients that benefit varies widely by cancer type and we currently lack biomarkers with which to reliably predict immunotherapy response^10^. Emerging evidence from clinical trials indicates higher response rates in those cancer types that typically display greater lymphocyte infiltration (e.g. melanoma, lung cancer, head and neck cancer) and that the tumour neoantigen repertoire (a function of somatic mutation load) is a key determinant^11–14^. These observations point to a model in which, within any given cancer type, there are ‘immune hot’ and ‘immune cold’ tumours. Immune hot tumours display greater cytotoxic T-lymphocyte (CTL) infiltration, and reactivation of these tumour-resident CTLs by checkpoint inhibition can result in dramatic tumour regression. Conversely, immune cold tumours display minimal CTL infiltrates and typically fail to respond to checkpoint modulation. If one could accurately identify likely responders for patient stratification and devise strategies by which to convert cold tumours to hot tumours, these would be major steps forward in realising the full clinical potential of cancer immunotherapy. Applying a novel method to estimate tumour composition from DNA methylation data, we set out to address these questions by identifying immune hot and cold tumours across a broad spectrum of cancer types. We aimed to understand the differences in the cellular composition of these tumours and to uncover any common genomic and transcriptomic alterations that are associated with immune responses or immune evasion.

Although flow cytometry of disaggregated tumour biopsies is commonly used for investigating cellular composition, this is often unfeasible for several reasons; difficulty in obtaining fresh tumour tissue; lack of defined markers for poorly characterised cell types (e.g. CAFs); and high cost of labour, reagents and equipment required for such analyses. Cellular disaggregation of collagen-rich tumours is also problematic, where cells are embedded in a dense extracellular matrix. To overcome these difficulties, multiple reference-free or reference-based methods have recently been developed to permit the in-silico deconvolution of complex cellular mixtures or to estimate tumour purity ^15–22^. For example, accurate deconvolution of complex cellular mixtures, including tumours, has recently been achieved by application of support vector regression modelling (CIBERSORT) to gene expression microarray data^21,23^. Notably, DNA methylation data are also suitable for deconvolution of tissue mixtures, although studies so far have focussed primarily on simple tissues such as blood, where cell type differences are a major confounder in Epigenome Wide Association Studies^17^.

Here we apply CIBERSORT-based deconvolution to genome-wide DNA methylation data from whole tumour tissue (hereafter referred to as ‘MethylCIBERSORT’) which permits accurate tumour cell deconvolution of both fresh and archival samples. Notably, unlike other available methods, MethylCIBERSORT estimates the tumour cell content (tumour purity) of a sample, in addition to performing deconvolution, thus providing absolute rather than relative estimates of infiltrating cell fractions. Initially focussing on head and neck cancer (HNSCC), a tumour type where we have previously demonstrated the prognostic significance of tumour-infiltrating lymphocytes (TILs), particularly in those cancers driven by human papillomavirus (HPV)^1,24^, we extended our technique to 21 further solid malignancies. As expected, the proportion of immune hot tumours varies widely by cancer type, but immune hot tumours are found even among those malignancies that typically display very little immune infiltration, such as pancreatic ductal adenocarcinoma or prostate adenocarcinoma. We leverage both DNA methylation and gene expression based deconvolution to reveal differences in the phenotype, from Th1*/*M1 pro-inflammatory in hot tumours, to Th2*/*M2 pro-fibrotic in cold tumours. Our genomic analysis reveals multiple copy number alterations enriched in cold tumours, including deletions in *PTEN* and amplifications in *MYC* and *EGFR*. We show that responses to PD1-blockade are associated with a transcriptional signature for hot tumours post-treatment, while the cold signature, and specifically a gene expression module we previously linked to increased aerobic glycolysis downstream of *EGFR* in HNSCC^25^, is enriched in non-responders. Importantly however, defining whether a tumour is hot or cold is not sufficient to accurately predict response to immune checkpoint blockade. By interrogating matched genomic data, we find that (presumably due to selective pressures imposed by the adaptive immune system during their evolution) treatment-naive hot tumours frequently display genomic alterations known to confer immunotherapy resistance. Our findings provide an explanation for the failure of immune checkpoint blockade in a subset of well-infiltrated tumours and identify genomic biomarkers for patient stratification. Building upon recent analyses of immune infiltrates and tumour gene expression profiles^20,23^ or molecular correlates of cytolytic activity^26^, we reveal fundamental, cross-cancer patterns of cellular infiltration and their relationships with genomic make-up with important implications for immunotherapy.

## Results

### Development and validation of methylation-based deconvolution using CIBERSORT

To develop a DNA methylation based deconvolution pipeline for application in tumours, we developed a custom R interface to develop basis matrices for use with CIBERSORT and generated a reference using fibroblasts and seven different immune cell types (see methods for details). We then evaluated the ability of our feature selection heuristic to accurately deconvolute mixtures of leukocytes using publicly available methylation data from mixtures of PBMCs with composition verified by flow-cytometry (gold standard). This showed extremely high correlation between the estimated and gold-standard fractions (Pearson’s R = 0.986, p < 2.2e-16, Figure 1A). We also carried out benchmarking against the performance of RNA-based CIBERSORT using the LM22 basis matrix against leukocyte mixtures of similar resolution originally profiled in Newman et al^21^. This revealed that MethylCIBERSORT estimates demonstrate higher correlations, both at the cell-type and the sample level (Figure 1B, C) and significantly lower absolute error (Figure 1D). Thus, methylation data coupled to CIBERSORT is highly accurate and may offer distinct advantages relative to expression-based CIBERSORT.

**Figure 1:**
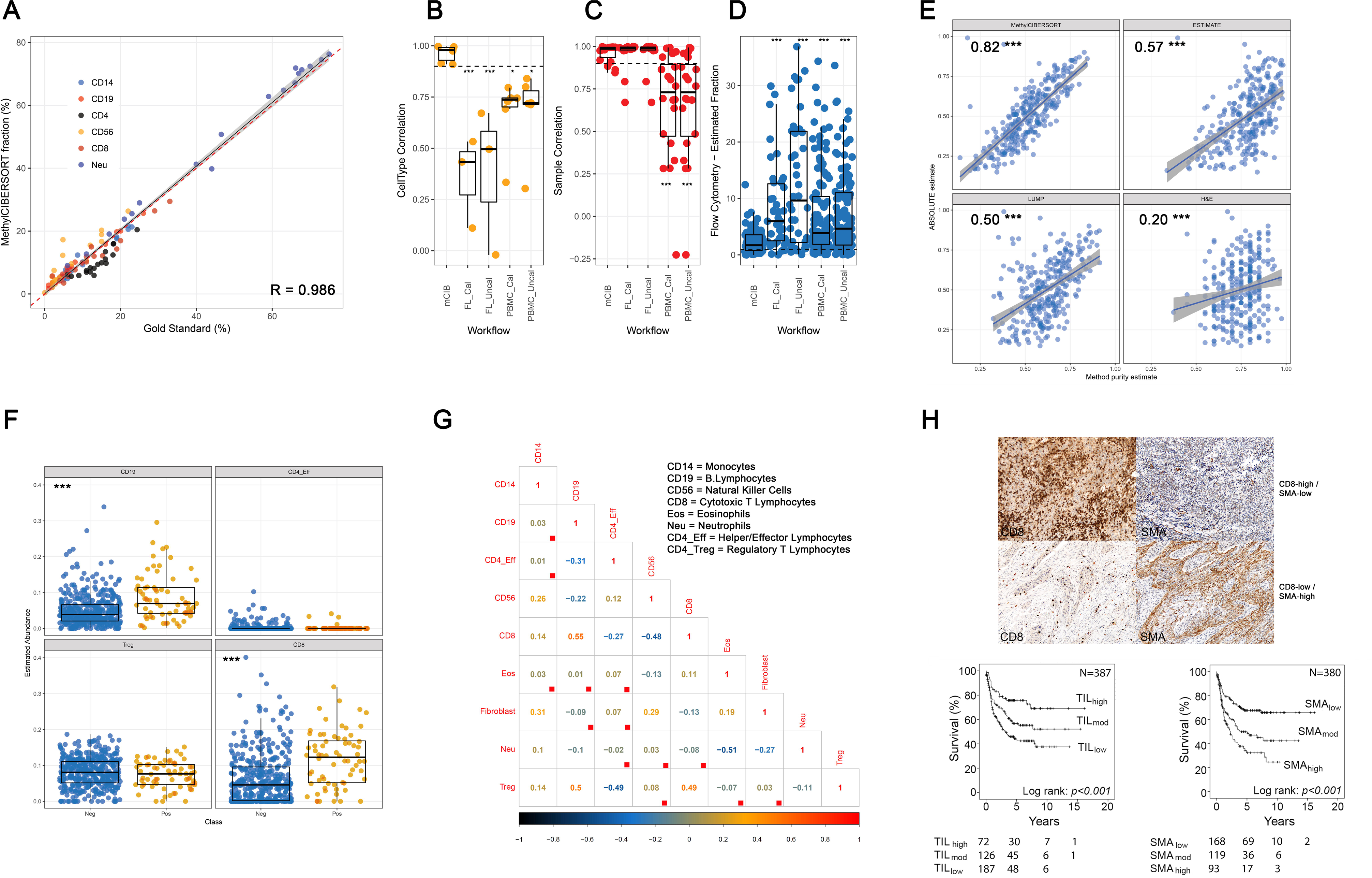
Validation of DNA methylation-based deconvolution for the analysis of tumour composition. **(a)** Correlation between MethylCIBERSORT fractions and flow cytometry for PBMC mixtures in independent data. **(b-d)** Boxplots showing comparisons between MethylCIBERSORT and flow cytometry versus Expression-CIBERSORT and flow cytometry in mixtures of similar complexity for correlations by cell type, correlations within samples, and finally absolute error. **(e)** Correlations between ABSOLUTE and MethylCIBERSORT versus other previously published purity estimation methods. **(f)** Validation of previously reported associations between CD8 T-cells and B-cells and HPV status by HPV status. **(g)** Correlation plot showing Spearman’s Rho between cell-types in HPV-HNSCC, red boxes indicate nonsignificance at q < 0.1. **(h)** IHC showing representative image of CD8 and SMA and Kaplan-Meier curves confirming the prognostic impact of TILs and fibroblasts in HPV-negative HNSCC.

To validate the method on real tumour samples, especially the tumour content in order to permit absolute quantification of tumour composition, we applied our pipeline to generate an HNSCC specific basis matrix and applied it to the set of 464 HNSCCs that have both RNA-sequencing and DNA methylation profiles available from The Cancer Genome Atlas (TCGA) project ^27^. Upon comparing cancer cell proportion (purity) estimates derived using MethylCIBERSORT with estimates derived from ABSOLUTE ^28^ (which jointly estimates purity and ploidy using mutation and copy number data) relative to other previously published methods of estimating purity (LUMP^29^ and ESTIMATE^30^) with data aggregated in^29^, MethylCIBERSORT displayed the highest correlation (R=0.82) and better concordance with ABSOLUTE than other methods (Figure 1E). Analysis of residuals (method estimate – ABSOLUTE estimate) suggested close concordance with ABSOLUTE estimates for MethylCIBERSORT, with larger deviations only seen when samples were of very high purity (>80%), while other methods tended to overestimate tumour cell content in samples of low purity (Figure S1A), resulting in statistically significant differences in distributions (FDR < 2.2e-16).

We also compared the mRNA expression of a panel of cellular lineage markers with MethylCIBERSORT estimates and found significant associations for multiple cell types (Figure S1B) even though they are derived from different samplings of the same tumour. Many of these marker genes demonstrated more variable expression in tumours with lower estimates of infiltrating cell fraction, suggesting that low coverage on either or both platforms (RNA-seq and methylation array) at the lower end of cellular abundance may result in poorer concordance. Taken together, these observations confirm that MethylCIBERSORT can accurately deconvolute the mixed cell populations in tumour samples using DNA methylation data.

### Detection of increased B- and T-lymphocyte infiltration in HPV-associated HNSCC

Having established the potential of MethylCIBERSORT to identify patterns of cellular infiltration in solid tumours, we tested its ability to detect the elevated TIL levels previously documented in HPV-driven (HPV+) HNSCC^1^. MethylCIBERSORT detected not only the increased TIL levels in HPV+ HNSCC compared with HPV-HNSCC (p= 2.167e-05, Wilcoxon’s Rank Sum Test) but more specifically attributed this to increased numbers of B (CD19+) and cytotoxic T (CD8+) lymphocytes (CTLs, Figure 1D), in agreement with observations made using other methods including immunohistochemistry and gene expression analysis^31^, potentially also helping to explain favourable prognosis displayed by this subgroup, independent of treatment modality^32–34^.

### Deep deconvolution highlights novel associations between infiltrating cell types and identifies two distinct patterns of infiltration in HNSCC

Next, we extended our analysis to HPV-negative (HPV-) HNSCC, a heterogeneous, anatomically-diverse group of tumours in which prognosis is typically much poorer than in HPV+ disease. Again, using TCGA data (available for 398 HPV-HNSCCs) we observed interesting relationships between multiple cell types, with 24*/*36 pairs of cell types showing significant correlations (Spearman’s rank correlations, FDR<0.01; Figure 1D). CTLs are associated with both CD14+ (monocytes */* macrophages */* myeloid-derived suppressor cells) and B-lymphocytes (Rho =0.14 and 0.55). CD4+*/*FoxP3- T-lymphocytes (CD4 + effector T-lymphocytes), meanwhile display inverse correlations with CTLs (R= - 0.27) and Tregs (R=−0.49). CD56+ Natural Killer (NK) cell abundance is also inversely correlated with CTLs (R=−0.48). Of note, CTLs are inversely correlated with fibroblast abundance (R=−0.16) and to validate this latter finding, we analysed data from two large studies in which these parameters had been quantified in HNSCC^1,7^. In a pooled analysis of these data, TIL content and SMA expression (a CAF marker) are inversely correlated (r=−0.322 and −0.344 for CD8 and CD3 IHC in the Ward (oropharyngeal SCC) cohort (Figure 1E); −0.4 and −0.424 for TIL scoring of H&E sections in the Ward (oropharyngeal SCC) and Marsh (oral SCC) cohorts respectively). They are also strongly prognostic (Figure 1F; p<0.001, Log Rank Test).

Given the complex nature of associations between different cell types, we performed consensus PAM clustering on the estimated cellular fractions to define subgroups by infiltration patterns. We derived two clusters (‘immune cold’ (C1) and ‘immune hot (C2)’ that show markedly different distributions of multiple cell types, most notably CTLs, CD4+ effector T-lymphocytes, CD19+ B-lymphocytes and NK cells, all of which are implicated in antitumour immunity (Figure 2A). Consistent with our previous observations, estimates of fibroblast content are higher in the immune cold group (mean fold change 1.31, FDR < 1.8e-6, Wilcoxon’s Rank Sum Test) and in multivariate Cox regression analysis, controlling for stage and age at diagnosis, membership of the Immune-cold group is associated with significantly shorter overall survival (Figure 2B; HR=1.42, CI=1.04–1.96, p=0.03). To explore the functional significance of our observations, we tested for associations between individual cellular fractions or immune cluster and a recently defined measure of local cytolytic activity based on the expression of Granzyme A and Perforin 1 (GZMA and PRF1; markers of activated T-cells)^26^. All infiltrating cell fractions display significant correlations with cytolytic activity, with CD8+ cells showing the maximum positive correlation (Figure 2C, FDR < 0.05, Spearman’s Rank Correlation). Accordingly, the immune hot cluster displays significantly higher cytolytic activity (Figure 2D, p = 2e-16, Wilcoxon’s Rank Sum test), and increased ratios of CTLs to Tregs (Figure 2E, p < 2e-16, Wilcoxon’s Rank Sum test); a metric that is prognostic in multiple settings^35–37^.

**Figure 2:**
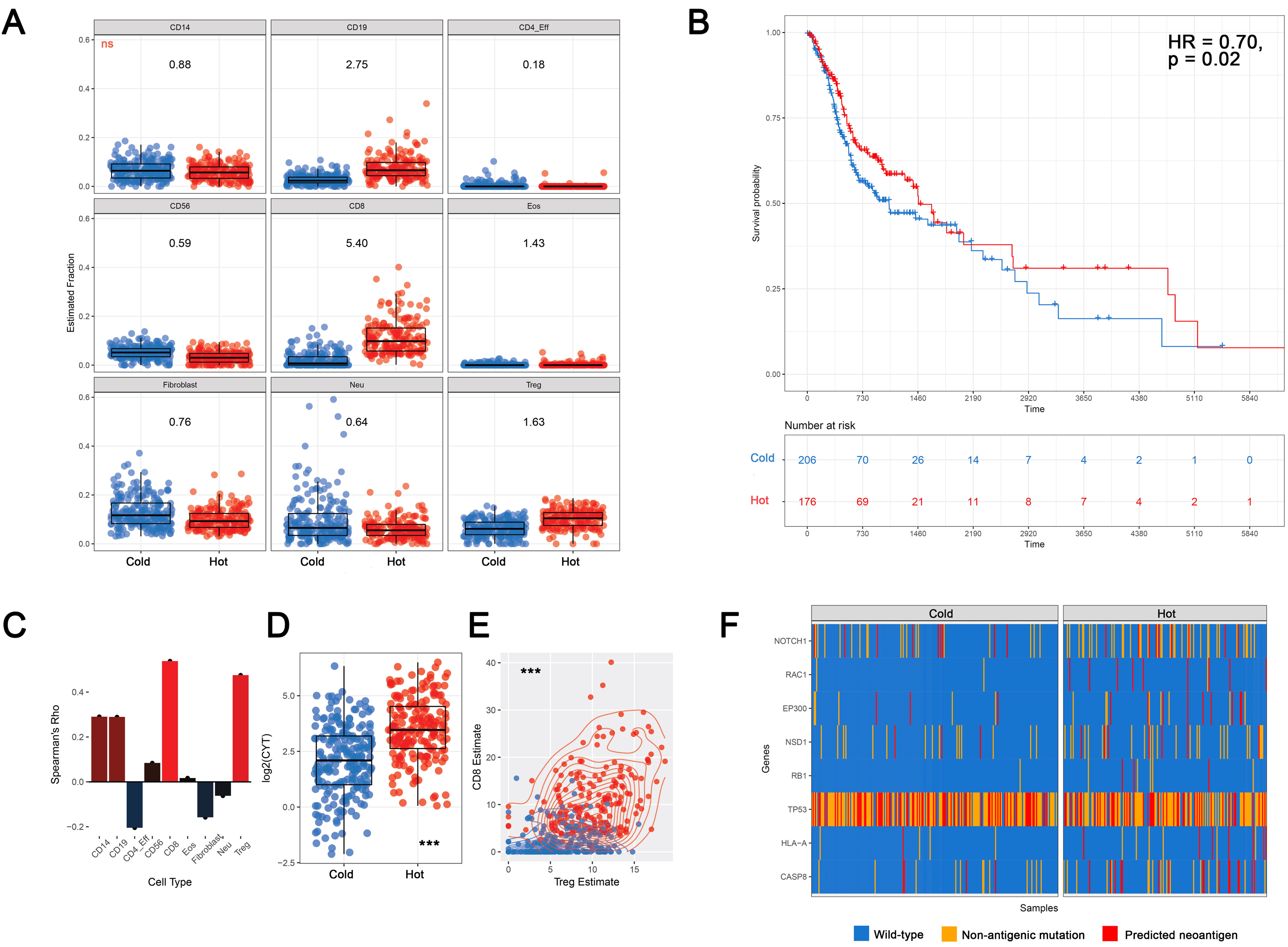
Classification of HNSCC into hot and cold tumour subgroups on the basis of immune cell infiltration patterns. **(a)** Boxplot of cell-types based on clustering HNSCC. **(b)** Kaplan-Meier curves of overall survival by HNSCC immune cluster. HR and p.values from multivariate Cox regression controlling for age and stage. **(c)** Bar-graph showing associations between cytolytic activity and cell types. **(d)** Cytolytic activity is elevated in immune-hot HNSCC. **(e)** CD8*/*Treg ratios by HNSCC Immune Cluster. **(f)** Mutations significantly associated with HNSCC immune cluster.

Together, our analyses suggest a tilting of the balance towards CTL activity in the microenvironment of tumours from the immune hot cluster that may explain the increased overall survival in this subgroup. Integrated analysis of the impact of different cell populations on cytolytic activity using linear modelling identified CD8+ (coef = 0.12, p < 2e-16), CD14+ (coef = 0.11, p <3.3e-8), Tregs (coef = 0.06, p < 0.003) and CD56+ (coef = 0.18, p < 8.8e-10) cell abundance as positive predictors and fibroblast abundance as a negative predictor (coef = −0.07, p <1.9e-10).

### Infiltrating cell patterns are associated with distinct transcriptional and proteomic profiles in HNSCC

Having established that these infiltrating cell patterns are of prognostic relevance in HNSCC we investigated if they are associated with different molecular profiles. Using limma-trend analysis, we identified 457 genes differentially expressed (DEGs) between the immune hot and immune cold clusters at a fold-change of greater than 2 (FDR<0.01, **Table S1**, genes highlighted in bold). Multiple DEGs are consistent with the MethylCIBERSORT-derived estimates of lymphocyte infiltration; *CD8A, ZAP70* and *CD3D* (CD8 lymphocyte markers), *CD79A* and *CD19* (B-lymphocyte markers), are all upregulated in the lymphocyte-enriched cluster, as are multiple chemokines and their receptors (*CCL5, CCR5, CXCR2, CXCR6, CCL19, CXCL11)*), immune checkpoint gene transcripts (*LAG3, PD1, IDO1, CTLA4*), immunosuppressive enzymes (*ADORA2A, IDO1*) and as expected, the cytolytic markers *PRF1* and *GZMA*. In extended analyses of all genes at FDR < 0.01 **(Table S1)**, multiple other genes, including the Class 1 MHC gene B2M (FC = 1.3), PD-1 ligand *CD274* (FC = 1.5) and *ACTA2*, which encodes SMA (FC = 0.71), are also differentially expressed between the two clusters, the latter validating fibroblast estimates from MethylCIBERSORT (Figure 2A). Of note, the increased expression of PD-1 ligand (PD-L1) the immune hot group suggests tumours of this subtype are more likely to respond to anti-PD1 checkpoint inhibition^38^.

Ingenuity Pathway Analysis further confirmed observations made using MethylCIBERSORT estimates, identifying differential regulation of multiple canonical pathways associated with immune function and inflammatory conditions **(Table S2)**, consistent with differential lymphocyte infiltration and activity. Diseases and functions ontology **(Table S3)** indicated that the top few pathways activated in the immune hot cluster were associated with leukocyte and lymphocyte migration. Upstream regulatory analysis implicated increased activation of the chromatin-modifying factors EHF and EZH2, and inhibition of Interferon-stimulated transcription mediated by IRF4, in the lymphocyte-rich tumours **(Table S4)**. In addition, OX40*/*OX40L signalling is predicted to be upregulated in immune cold tumours **(Table S2)**. OX40 is a co-stimulatory molecule expressed on T-lymphocytes and is one of a number of targets currently under early clinical investigation for immune checkpoint modulation therapy^39^. OX40 signalling opposes differentiation of CD4+ cells into Tregs and antagonizes Treg function, potentially explaining its reduction in Treg-rich immune hot tumours^40^. Finally, analysis of RPPA data identified 7 differentially abundant (FDR<0.1) proteins or phospho-proteins **(Table S5)**. Higher levels of cleaved Caspase 7 (FC=1.46) in the immune hot subgroup indicates increased apoptosis, whereas Fibronectin and PAI1 upregulation in immune cold tumours indicate a distinct pattern of TGFβ-driven extracellular matrix remodelling in what may be a CAF-linked phenomenon.

### Distinct mutations are associated with HNSCC immune cluster

Having established that immune cluster is associated with distinct transcriptional patterns, we then sought to identify individual mutations in driver genes (MutSig CV ^41^ q.value < 0.01) associated with immune cluster using Negative Binomial regression. This identified *CASP8, NSD1, NOTCH1, EP300, HLA-A, RAC1, and RB1* as significantly more mutated in immune hot HNSCC and *TP53* to be less mutated (Figure 2F).

*CASP8* mutations are implicated in subverting apoptosis induced by lymphocytes; they are enriched in tumours with high immune cytolytic activity and likely reflect an increased selective pressure exerted by the presence of adaptive immune cells^26,42^. Fas-ligand *(FASLG)*, an upstream activator of pro-apoptotic signalling through Caspase 8^43^ is also upregulated in the immune hot tumours, further highlighting the importance of this pathway **(Table S1)**. Identification of this lymphocyte-rich, good prognosis group displaying *CASP8* mutations and a relative lack of *TP53* mutations is striking, since TCGA previously identified a subset of good-prognosis oral cavity tumours bearing the same genomic hallmarks, which were reported to co-occur with *HRAS* mutations^27^.

Neoantigen burden has previously been identified as a predictor of anti-tumour immune responses^26,44,45^ and consistent with this, we identified significantly higher neoantigen burdens in the immune hot cluster (OR = 1.54, p < 1.8e-6, Negative Binomial GLM) and a smaller increase in overall mutational burden (OR = 1.23, p = 0.008). Strikingly, the differentially mutated driver genes by themselves tended to encode predicted neoantigens significantly more often in the Immune-hot cancers (OR = 1.44, p = 0.01, Fisher’s Exact Test). Moreover, in 16 tumours from the immune hot cluster versus 4 in the immune cold cluster, *CASP8* mutations themselves encoded at least one neoantigenic peptide (Figure 2F), demonstrating the existence of mutations that both contribute to the development of a potential selective constraint, and serve as an adaptive mechanism to evade it.

Our findings provide new insight into the *CASP8*-mutant */ TP53*-WT */ HRAS*-mutant prognostic subgroup identified by TCGA and suggest increased CTL infiltration as a potential mechanism to explain the improved outcomes seen in these tumours. The ability of our approach to rediscover a prognostic subgroup previously defined by genomic profiles using an independent approach highlights the potential of DNA methylation based immune cell fraction deconvolution. We then sought to use our inferences in HNSCC as a tool for investigating the immune microenvironment pan-cancer, permitting analyses where variation between tumour types could be accounted for to produce a comprehensive molecular portrait of cancer immunosurveillance.

### HNSCC-derived immune clusters are reproducible across tumour types

To examine whether the relationships between tumour composition, genomic alterations and clinical behaviour we observed in HNSCC are generally applicable, we derived cancer-type specific basis matrices) and conducted deconvolution on 18 further tumour types for which cancer cell line methylation data have recently been published^46^. For 9 of these we were able to compare our predictions of tumour purity with ABSOLUTE estimates and observed strong correlations and significantly lower error margins compared to LUMP and ESTIMATE(Figure S2A, **S2B)**. Further, we observed a robust preservation of positive correlations between MethylCIBERSORT and marker expression pan-cancer (Figure S2C), again with the caveat that the samples were taken from different aliquots of the tumour. Taken together, these findings attest to the general pan-cancer applicability of MethylCIBERSORT. An important potential advantage of DNA-methylation over gene expression-based deconvolution methods is the ease with which accurate DNA methylation profiles can be obtained from formalin-fixed, paraffin-embedded (FFPE) samples ^47–49^. We therefore compared estimates pertaining to fresh frozen and matched FFPE samples (n = 21 from 3 tumours)^50^ and recorded very high correlations, indicating our method is applicable also to archival material (Figure S2D).

We then trained an elastic-net classifier using 5-fold cross-validation for tuning on the HNSCC cellular abundance data, returning highly accurate recapitulation of clustering (Kappa = 0.9), and predicted immune cluster membership for the validation set of 7269 samples representing 21 further tumour types from TCGA (Figure 3A). As expected, we observed strong enrichment for CTLs, Tregs, and B-lymphocytes in immune hot tumours pan-cancer, while CD4-effectors, NK cells, eosinophils and CAFs were enriched in immune cold tumours (Figure 3B). Different tumour types also display markedly varying degrees of lymphocyte infiltration, with the majority of pancreatic ductal adenocarcinomas, colorectal, thyroid, uterine corpus endometrial, kidney, prostate, hepatocellular cancers and sarcomas belonging to the immune cold cluster (Figure 3A). We again observed increased CTL:Treg ratios in immune hot tumours (Figure 3C) and similar relationships between tumour composition and CYT to those seen in HNSCC.

**Figure 3:**
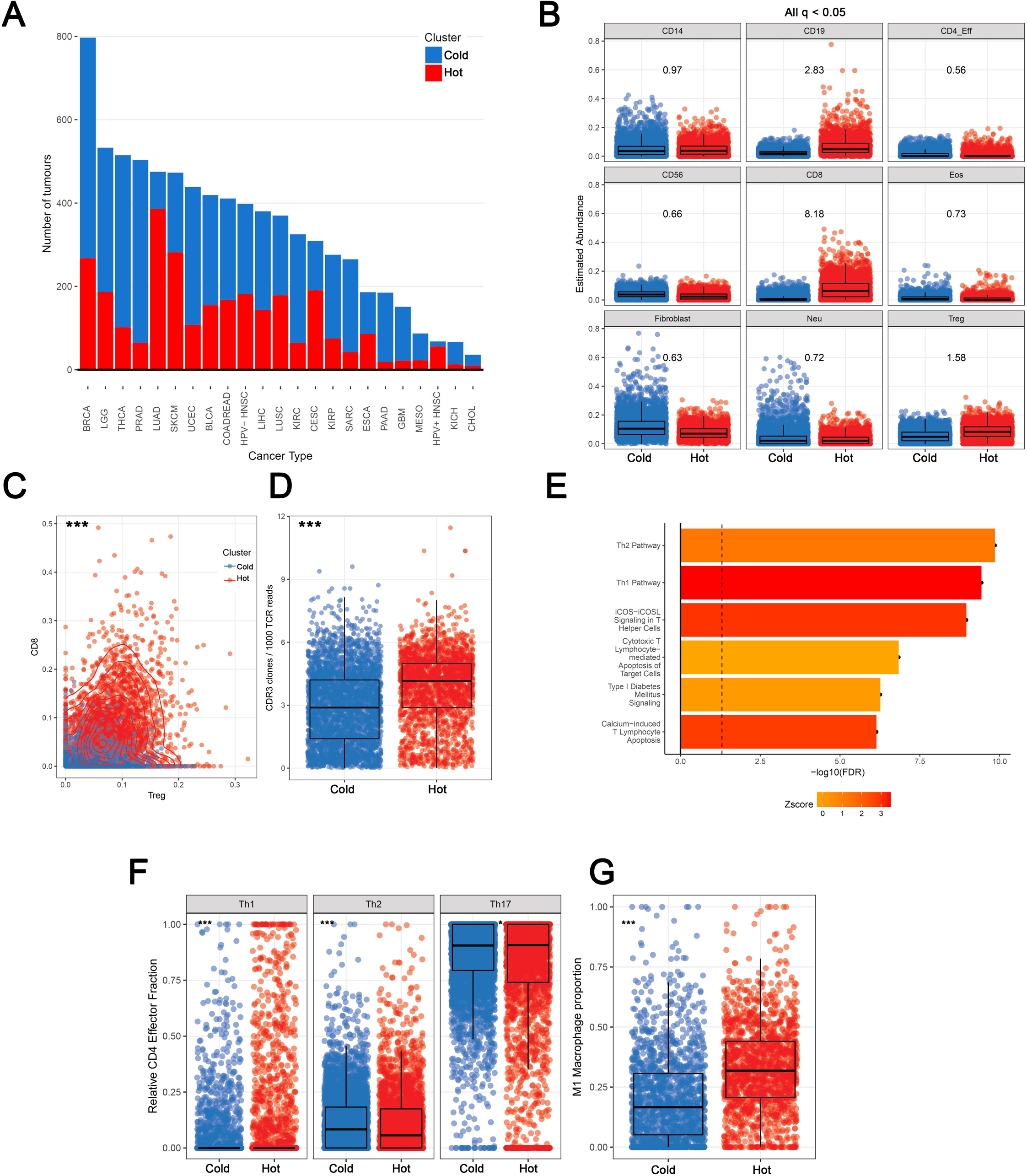
Identification and characterization of hot and cold tumours pan-cancer. **(a)** Barplot of distribution of Immune-hot and cold tumours across TCGA. Cancers known to respond favourably to checkpoint blockade, such as lung cancer and melanomas, show high fractions of hot tumours. **(b)** Boxplot of cell-type estimates by immune cluster. All at q < 0.05. Numbers represent mean fold changes. **(c)** CD8:Treg ratio is elevated in hot tumours pan-cancer. **(d)** Increased breadth of TCR sequences in Immune Hot tumours. **(e)** Results of IPA canonical pathway analysis comparing hot and cold tumours pan-cancer after adjusting for tumour type. **(f)** Transcriptional deconvolution by Expression-based CIBERSORT shows immune cluster is associated with distinct CD4 polarisation and **(g)** macrophage polarisation.

### Immune hot tumours display greater evidence of immunoediting and are marked by Th1/M1 responses

To further determine if the immune infiltrate was active in these tumours, we assayed immunoediting by testing for reductions from the expected ratio (as previously defined by Rooney et al^26^) of observed neoantigens to total nonsilent mutations per tumour and adapted this approach to derive the estimated number of neoepitopes lost through immune editing while controlling for tumour type. Accordingly, we found significant enrichment for editing in immune hot tumours compared to immune cold tumours (OR = 1.23, p = 0.008, Negative Binomial GLM). Additionally, upon integration with T-cell receptor (TCR) repertoire data from Li et al^51^, we found more diversity (Number of TCR clones */* Total number of TCR reads) in the immune hot tumours (Figure 3D, **p < 2.2e-16)**, suggesting that broader immune responses may underlie the greater depletion of neoantigens in this group.

Given the evidence for divergent infiltration patterns and activity between the immune clusters across cancer types, we then investigated the determinants of this response by identifying differentially expressed genes after adjusting for tumour type. We identified 275 genes at FDR < 0.01, 2FC and in pathway analysis, the top pathways were significantly associated with Th1 vs Th2 responses (Figure 3E, **Table S6)**. Multiple Th1 cytokines and downstream targets were overexpressed in hot tumours (*IFNG, CCL4, CCL5, CXCL9, CXCL10*), along with costimulatory and coinhibitory receptors, suggesting these tumours were marked by a state of lymphocyte activation and counter-responses thereto. We next scored proinflammatory (Th1, Th17) and suppressive (Th2)) CD4+ cell populations using RNA-seq reference profiles from purified cells to derive relative estimates using CIBERSORT^21^. Consistent with our inferences from pathway analysis, we found enrichment for Th1 and Th17 cells in immune hot and Th2 cells in immune cold cancers (Figure 3F), translating to markedly elevated Th1*/*Th2 ratios in hot tumours (p < 2.2e-16, Wilcoxon’s Rank Sum Test), with smaller but significant increases in Th17*/*Th2 ratios (p = 1.2e-5, Wilcoxon’s Rank Sum Test). Importantly, T-helper 2 (Th2) cells have been linked to poor prognosis in multiple studies, while Th1 cells are associated with good prognosis and aiding CTL responses^52^.

We also used expression-based CIBERSORT to derive estimates for different myeloid cell populations (n = 2346 tumours at deconvolution P < 0.05), and identified substantially higher fractions of M1 relative to M2 macrophages in hot tumours (p = 2.2e-16, Wilcoxon’s Rank Sum Test, Figure 3G). Notably, M2-like polarisation is associated not only with Th2 immune responses but also with immune-suppressive Myeloid Derived Suppressor Cells (MDSCs)^53^. Taken together, our analyses implicate Th1*/*Th17cytokine signalling programmes as responsible for establishing an immune-hot state and define MDSCs and Th2 programmes as targets for efforts to switch immune cold tumours to an immune hot state.

### Transcriptional correlates of immune clusters predict responses to checkpoint blockade and suggest TCGA hot tumour transcriptomes are linked to active antitumour immunity

We reasoned that if the signature for immune hot tumours represented active immunity, it would be applicable to the prediction of immunotherapy responses and evaluated this hypothesis using tumour gene expression data from three melanoma cohorts: post-sequential aCTLA4 and aPD1 treatment^54,55^; pre-aCTLA4 treatment^14^ and post-aPD1 (Nivolumab) treatment^56^. Analysis of these transcriptional patterns indicated differential expression between responders and non-responders (Figure 4A), and accordingly, ssGSEA scores for the hot transcriptional signature showed significant enrichment in responders for the latter two datasets (Figure 4B, C). Moreover, a similar association emerged from comparing the probability of response to hot*/*cold class prediction, inferred using a logistic regression fit on TCGA hot*/*cold transcriptional signature ssGSEA scores (Figure 4D). Finally, we evaluated the ability of the hot-signature to stratify patients by response relative to mutational load and Class I neoepitope burden using elastic nets coupled to cross-validation for each dataset (Figure 4E). This showed that in the post treatment data, the immune-hot signature outperforms neoantigen and mutational burdens respectively, and for pretreatment aCTLA4 data, performs similarly to mutational burden and neoantigen burden. Notably, however, even in the post-treatment setting, the presence of the hot signature does not translate to a guaranteed response, indicating potential prior selection for genetic and epigenetic alterations that confer resistance to T-cell mediated killing in otherwise immunogenic tumours

**Figure 4:**
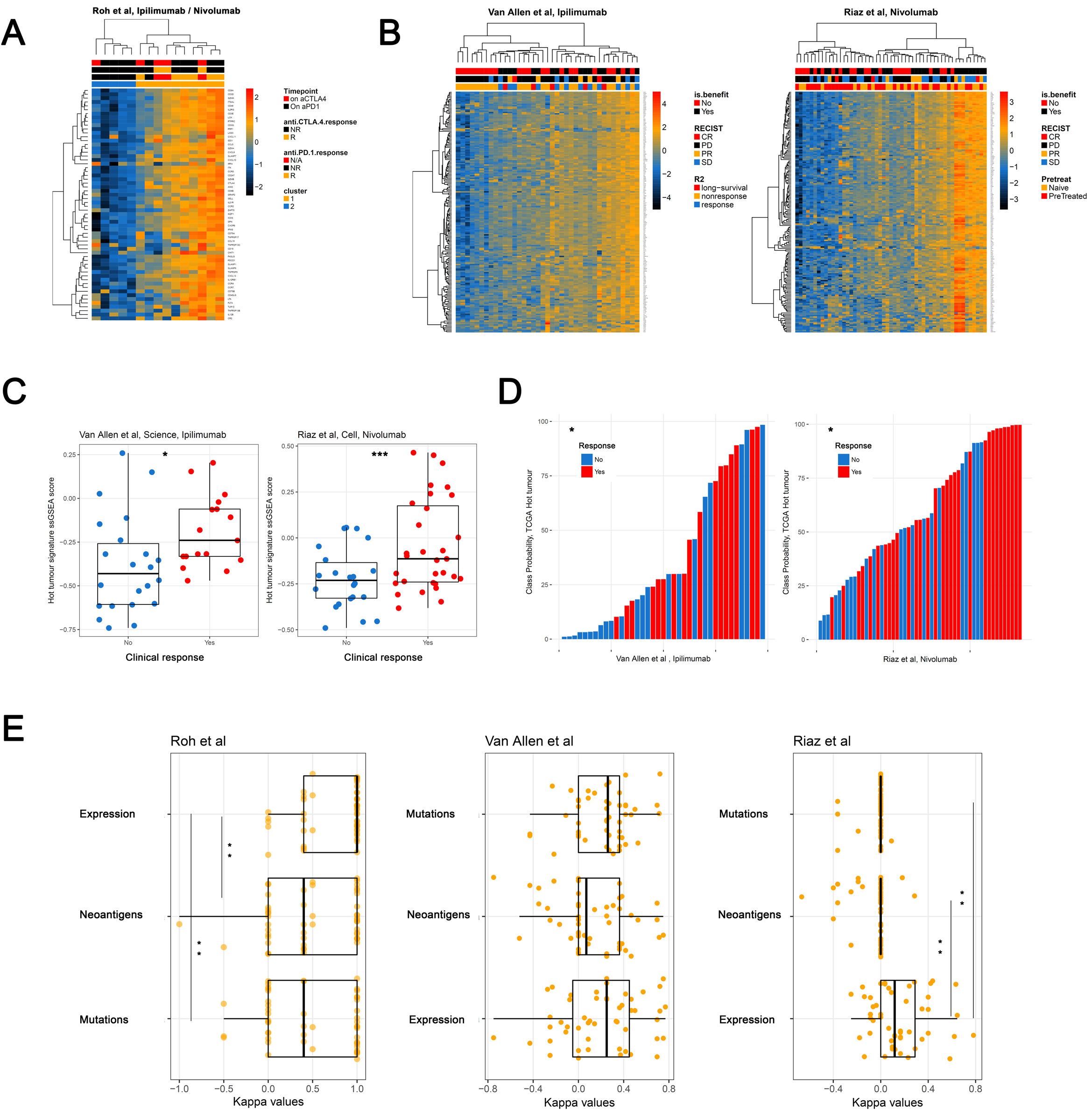
The immune-hot signature is associated with response to immune checkpoint blockade in melanoma. **(a)** heatmap showing expression of the hot-tumour transcriptional signature in Nanostring data from posttreatment biopsies of immunotherapy patients. **(b)** heatmaps showing the same signature in RNAseq data of aCTLA4 (pre-treatment) and aPD1 (post-treatment) respectively. **(c)** boxplots highlighting significant differences in ssGSEA scores for the hot-tumour transcriptional signature in the datasets featured in (b). **(d)** barplots display similarity to TCGA hot and cold tumours based on logistic regression class probabilities from a model fit to TCGA data, which are associated with response. **(e)** boxplots showing Kappa values from cross-validation for models examining the performance of the Immune-hot signature, Class I neoepitope burden, and finally mutational load on immunotherapy response classification.

Taken together, these findings establish the fundamental similarity between cancers responsive to immunotherapy and immune-hot tumours across a wide range of cancer types, and suggest that genomic features that determine immunotherapy response should also be enriched in immune hot TCGA tumours. We reasoned that the heterogeneity in responses to checkpoint blockade among hot tumours might be driven by intrinsic resistance to T-cell mediated destruction due to pre-existing genomic alterations within the tumour cells. We set out to test this hypothesis by constructing a pan-cancer catalogue of genomic alterations enriched in hot tumours, with the additional aim of finding those alterations enriched in cold tumours which might drive lymphocyte exclusion or reduce tumour immunogenicity.

### Genomic features of hot and cold tumours

Consistent with our observations in HNSCC, and with estimates from gene expression-based deconvolution^20^, immune hot tumours harboured higher overall mutation loads (OR = 1.4, p = 1.64e-31, negative binomial GLM controlling for cancer type) and more predicted neoantigens than immune cold tumours (Figure 4A).

Given the recent finding that in addition to the presence of neoantigens, their clonality (i.e. presence in all tumour cells as opposed to minor subclones) is associated with prognosis and response to Pembrolizumab in lung adenocarcinoma^12^, we analysed immune microenvironment composition as a function of neoantigen clonality (as denoted by The Cancer Immunome Atlas^57^). We found that the abundance of both CTLs and Tregs is correlated with clonal neoantigen load pan-cancer (Figure 4B), while the relationship is much weaker when subclonal neoantigens are considered. CD4+*/*FOXP3-effector lymphocytes display a striking inverse correlation with clonal neoantigens (Figure 4C). Consistent with our earlier observation they are enriched in CTL */* Treg low immune cold tumours CAFs are inversely correlated with both clonal and subclonal neoantigen loads. Immune hot tumours display a significantly higher clonal neoantigen burden (OR = 1.236, p < 2.2e-16, Negative Binomial GLM) as well as a skew in the neoantigen burden towards clonal neoantigens after adjusting for tumour type (OR = 1.03, p = 1.6e-5). These findings provide the first evidence for a direct link between Class I MHC clonal neoantigen burden and patterns of TIL abundance and help to explain the observations of McGranahan and colleagues, that high clonal neoantigen burden predicts favourable response to immune checkpoint modulation using Pembrolizumab^12^. While it is unclear why clonal neoantigens should elicit a proinflammatory Th1 response, some previous work has suggested that the antigen dose may determine the nature of the subsequent immune response^58^, or alternatively, clonal neoantigens may simply have been subjected to immunosurveillance for far longer, and this is borne out by improved immunosurveillance in mouse models when mismatch repair deficiency is induced clonally instead of subclonally^59^.

Finally, we examined if the genomic features associated with immune cluster were also reproducible across cancer types, performing adjusted binomial regressions to estimate cluster association after controlling for tumour type for genes previously implicated as pan-cancer drivers based on signatures of positive selection^60^ and recorded 114 hits at FDR < 0.1 (Table S7). Interestingly, these putative drivers of hot tumours were significantly enriched (OR = 8.43, p < 0.002, Fisher’s Exact Test) in a list of genes demonstrated to confer resistance to CD8 T-cell mediated killing in a recent CRISPR-Cas9 screen ^61^. These functionally-verified immune-resistance genes included components of the MHC Class I complex such as *B2M* and *HLA-A*, apoptosis pathway genes such as *CASP8*, and *JAK1*, which is required for interferon mediated signalling that in turn is associated with resistance to checkpoint blockade^62–64^. We also show the recently-identified immunotherapy sensitizers (*ARID2, BRD7*^65^) to be disproportionately mutated in hot tumours, together with tumour suppressors such as *RB1* ^66^, *TP53* ^67^, *LATS2* ^68^, *ATRX* ^69^ *and SETD2* ^70^, all of which have been associated directly or indirectly with interferon responses, additional epigenetic regulators (*KMT2A, TET2, IDH1, NSD1, KDM6A, KMT2B, BCOR, and NCOR*), and finally, multiple DNA-damage associated proteins, including *BRCA1, BAP, TOP2A* and *CDK12*. Taken together, these point to a model where mutations in certain genes render tumours hot as a consequence (and therefore susceptible to checkpoint blockade), or may enable tumours to survive in a hot tumour microenvironment, potentially also bestowing resistance to checkpoint blockade. We sought to test this model by linking immune-resistance mutations to lack of immune checkpoint blockade response in hot tumours and although we observed a trend, (odds ratio= 0.26), the number of treated tumours with sequence data available is currently too low (41 hot tumours across four studies) to gain a definitive answer.

To complement these analyses, we also carried out copy number analyses, calling copy numbers across 11,000 tumours and testing for differential association of peaks with immune cluster after adjusting for tumour type for the subset with Immune Cluster Assignment available. This led to the identification of multiple events that occurred at different frequencies between cold and hot tumours (FDR < 0.1, Figure 4B). Of these, prominent examples included amplifications targeting the epidermal growth factor receptor (*EGFR*) (7p11.2) and *MYC* (8q24.3) and deletions at 10q23.31, encompassing the *PTEN* tumour suppressor gene in cold tumours and *JAK2* (9p24.1) amplifications in hot tumours. Some of these candidates already have known associations with immune evasion; *MYC* has been linked to an immune evasion phenotype that is amenable to targeting through gene-body demethylation ^71^ and *PTEN* deletion has recently been linked to failure of immunotherapy and decreased cytotoxic T-lymphocyte infiltration in patients and in a mouse model of melanoma^55,72,73^. Among the genomic alterations we identify (for full list of predicted driver events see **Table S8**), it is likely that some establish, while others are selected for, in different immune microenvironments. In either case, alongside *PTEN* deletion, these alterations warrant further investigation as candidate genomic markers for response to immune checkpoint blockade. The enrichment of *EGFR* and *MYC* amplification, together with *PTEN* deletion in cold tumours pan-cancer was striking given the co-expression module linked to increased tumour cell glycolysis and immune evasion in HNSCC, which includes *EGFR* and in which pathway analysis also predicts increased c-MYC and mTORC1 activity^25^. A similar relationship has been observed in triple negative breast cancer^74^ and we therefore investigated this relationship further, initially interrogating the link between *EGFR* protein levels and TILs in two HNSCC cohorts (n=518)^1,7^ we found that samples classified as *EGFR* high and moderate were significantly more likely to be TIL low than *EGFR* low cancers after accounting for anatomic site and HPV status (Figure 5E, p < 0.05 and 0.01 for *EGFR* moderate and high cancers, Logistic Regression). The positive correlation between *EGFR* levels (which are themselves correlated with *EGFR* phosphorylation (activation), Figure S3) and the glycolytic signature is maintained across TCGA when matched RPPA profiles and RNA-seq data are compared (Fig 5E). Notably, high levels of the glycolytic signature are present in progressing melanomas after PD-1 blockade (Fig 5F, p = 0.06, t-test, p = 0.02 when excluding stable disease) and are inversely associated with expression of the immune-hot transcriptional signature (Rho = −0.44) we associated with response (Fig 5G).

**Figure 5:**
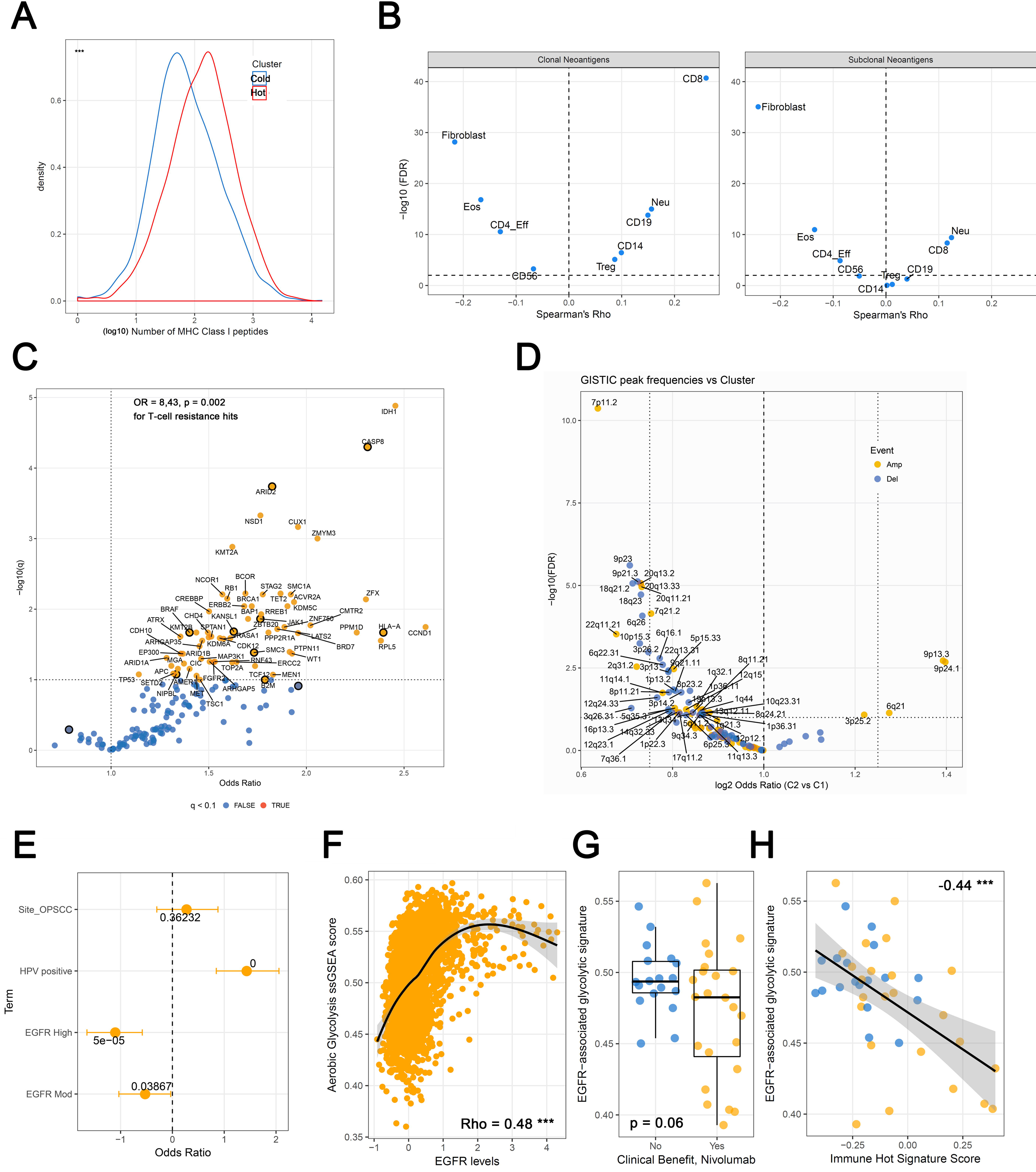
Genomic features of hot and cold tumours. **(a)** Density plots showing differences in neoantigen burden by immune cluster pan-cancer. P.value from negative binomial regression that accounts for tumour type. **(b)** Clonal neoantigens and subclonal neoantigens are correlated with different infiltration profiles. Volcanoplot shows Spearman’s Rho on the X-axis and –log10(FDR) on the y-axis. **(c)** Volcanoplot showing results of binomial regression testing for associations between Immune-hot cancers and mutation frequencies in candidate cancer driver genes. Those genes implicated in resistance to T-cell mediated destruction are highlighted in orange. **(d)** Volcanoplot showing associations between GISTIC candidate driver copy number peaks and immune cluster. **(e)** Plot showing results of logistic regression in a cohort of HNSCCs where the probability of being classified TIL-high was regressed against anatomic subsite, *EGFR* IHC (low*/*moderate*/*high) and HPV status. **(f)** Correlation between glycolytic coexpression signature ssGSEA scores and *EGFR* levels by RPPA. **(g)** Association of glycolytic signature post-Nivolumab with response and **(h)** Inverse correlation between the glycolytic signature and the immune-hot expression signature, Spearman’s correlation has been plotted.

## Discussion

Of the methods developed to deconvolve cell mixtures into multiple cell types from methylation data, none have been comprehensively employed across cancer types. Methods such as LUMP and the Leukocyte Methylation Score estimate only the overall leukocyte fraction, while methods based on expression data either produce relative estimates of abundance within the immune fraction or enrichment scores(CIBERSORT, TIMER^20^) or perform low resolution deconvolution (ESTIMATE)^21,29,75,76^. Combining methylation-based feature selection from both stromal and cancer cells with the robust performance of CIBERSORT previously displayed on gene expression microarray data^21^ allowed us to derive estimates for different infiltrating cell populations as a fraction of the overall sample.

While approaches using RNA-sequencing or other transcriptional profiling, such as the construction of an index of cytolytic activity, have been useful in predicting immunotherapy response^77^ and in identifying the role of mutations in genes like *CASP8* in immune evasion^26^, the deeper level of deconvolution made feasible using DNA methylation data allows the roles of distinct cellular subsets and their interdependencies to be dissected. Here, by applying the method to HNSCC, which is marked by a great degree of clinical heterogeneity, we identified lymphocyte-rich and stromal-rich prognostic subgroups consistent with those discovered previously using a variety of independent methods ^1,7,31,78–81^ and derived novel insights into the microenvironmental alterations that modulate prognosis. In the process, we showed that our scheme for classifying cancers correlates with well-established immune metrics such as cytolytic activity, neoantigen*/*mutational load, and CTL:Treg ratios, and then demonstrated that tumours similar to the HNSCC Immune-hot subgroup, which by association exist in varying fractions across the vast majority of cancer types, and the congruence of our classification with the aforementioned metrics is maintained throughout, and translates to broader TCR responses which in turn correspond to greater depletion of neoantigens.

We then demonstrated that the transcriptional correlates of our classification scheme are indicative of active antitumour immunity through Th1 responses and reinforced this theme through integration with RNA-based CIBERSORT that indicated differences in CD4 and macrophage polarisation. We went on to show that the hot-tumour transcriptional programme, if induced upon checkpoint blockade, is strongly associated with response, indicating that it represents active antitumour immunity and establishing the TCGA as a resource to study genomic alterations that may associate with checkpoint blockade resistance*/*sensitivity. Upon extending our analysis pan-cancer, we made several observations that give novel insight into the interplay between tumour genomics and the immune microenvironment. Genomic analysis identified significant enrichment for events that confer resistance to T-cell mediated destruction in hot tumours as well as potential sensitisers. Our copy number analysis revealed that *PTEN* deletion, *MYC* amplification and *EGFR* amplification are associated with immune depletion.

All of these mediate increased glycolysis, which we have previously linked to immune evasion^82^. Our finding that *PTEN* deletion is associated with poor CTL infiltration in this pan-cancer cohort adds substantial support and mechanistic rationale for its proposed role as a determinant of response to immune checkpoint blockade. Taken together with our identification of *EGFR* and *MYC* amplification in cold tumours, our analysis suggests that pharmacological inhibition of EGFR/mTORC1*/*MYC-driven glycolysis could be an effective means by which to ‘warm-up’ these tumours and potentially enhance responses to immune checkpoint blockade. Additionally, our analysis of neoantigen clonality and immune infiltration patterns adds mechanistic insight to the value of clonal neoantigen burden in predicting response to immune checkpoint blockade^12^. In particular, we show that clonal neoantigens are associated with infiltration of CTLs and Tregs, while Th2 cells and CAFs are enriched in tumours with lower clonal neoantigen loads. These findings support recent evidence suggesting that differentiation of naïve CD4+ T-lymphocytes into Tregs occurs within tumours^83^, since a microenvironment favouring differentiation into Tregs would likely be selected for in tumours with increased neoantigens and more infiltrating CTLs. Why these relationships between neoantigen loads and T-lymphocytes are apparent only when one considers clonal neoantigens is an intriguing question. It could be that since many subclonal neoantigens are expressed by a small minority of cells within the tumour, these evade effective presentation to the immune system. Indeed, in a previous study by several of the authors, it was possible to isolate T-lymphocytes reactive against clonal but not subclonal neoantigens from lung cancer patients^12^. Our data suggest that this is due to a relative paucity of CTLs in tumours with low clonal neoantigen loads and that this is true across a wide range of cancer types. In summary, the development of a stand-alone method to estimate both tumour purity and stromal composition from DNA methylation data has allowed us to make a number of novel insights that shed light on potential biomarkers for immunotherapy response and the way in which tumour genomes influence, and are shaped by, the immune microenvironment. Finally, the association between neoantigen clonality, the probability of being immunologically hot, and enrichment for known mediators of immune evasion*/*resistance in hot tumours before treatment with any checkpoint blockade also raises profound questions about the use of immunotherapy; the features that render tumours susceptible to immune destruction are likely to have existed far back into the evolutionary history of these tumours, increasing the likelihood that natural selection will have already produced genomic alterations that doom eventual checkpoint blockade to failure. Thus, we can stratify immune-hot tumours into two groups: those without aberrations in immune-resistance genes that we may expect to respond to checkpoint blockade, and those with resistance mutations, in which these either these aberrant pathways will likely have to be co-targeted, or alternative therapies may be more suitable. Finally, the lack of immune-resistance mutations in cold tumours (presumably due to the absence of a selection pressure for them) suggests that if we can induce lymphocyte infiltration (e.g. by targeting glycolysis or CAFs^84^), we may improve the effectiveness of checkpoint blockade across a broader range of patients.

## Methods

### Code Availability

All scripts and functions developed for our method will be made freely available in an R-package upon publication of this manuscript. R markdowns for analysis code will be available by request upon publication.

### Development of a methylation signature for in-silico deconvolution

#### Dataset Assembly and Preprocessing

Raw data were obtained in the form of IDAT files from the following sources (the number of samples from which each profile was derived is shown in parentheses): Granulocytes (12), CD8+ (cytotoxic T-lymphocytes) (6), CD19+ (B-lymphocytes) (6), CD56+ (Natural Killer cells) (6), CD14+ (monocyte lineage) (6), Eosinophils (6) were from the Blood.450k Bioconductor package^85^. CD4+ cells were removed from the Blood.450k dataset and CD4+ T-cells from the Zhang dataset^86^ (data kindly provided by Dr Alicia Oshlack) were further divided into FOXP3+ (Tregs) (4) or FOXP3-(6) groups. Fibroblast profiles (4) were from the Gene Expression Omnibus (GSE74877). Neutrophils are the most abundant subset of granulocytes and these samples were therefore aggregated into a single category for further analysis. To generate a DNA methylation signature for cancer cells, we used 450k methylation profiles we previously obtained from a series of 6 HNSCC cell lines: UM-SCC47; 93VU147T; UPCI:SCC090; PCI-30; UPCI:SCC036 and UPCI:SCC003 (GSE38270, described in^87^) and additionally those from Iorio et al (GSE68379).

The files were parsed into R using the *minfi*^88^ Bioconductor package and were normalised using single sample Noob as implemented in *minfi*.

#### Derivation of signature features

A custom limma based wrapper function was used to fit a series of linear models for all pairwise comparisons between candidate cell types. Features from this set of analyses were then restricted to MVPs that showed a median beta-value difference of 0.25 at an FDR of 0.01 for that fit or less, with a maximum of 100 MVPs per pairwise comparison. Finally, for use with CIBERSORT, data were transformed from beta values (bound between 0 and 1) to percentages (0 – 100). Type-wise means were estimated for each probe and cell type and the matrix exported for upload to CIBERSORT.

#### Benchmarking using PBMC mixtures

We applied the feature selection pipeline to the matrix of stromal cells that we assembled and then tested performance against 450k profiles of PBMC mixtures with flow-cytometry gold standards. We also applied *LM22* (Expression-based CIBERSORT) to datasets consisting of PBMC samples and Follicular Lymphoma biopsies originally evaluated in CIBERSORT^21^. Wilcoxon’s rank sum tests were used to test for differences in correlations with flow-cytometry for cell types and samples, and absolute errors between flow-cytometry and deconvolution estimate.

For the Expression CIBERSORT estimates, we performed comparisons against both calibrated (i.e, enforcing a sum-to-one constraint as reflected in the flow cytometry) and uncalibrated (straight estimates of cell fractions from CIBERSORT) estimates.

#### Running Deconvolution Experiments on HNSCC using CIBERSORT

Data for 464 methylation profiled TCGA HNSCC samples were downloaded in the form of raw IDAT files for the 450k array from the TCGA data. Data were normalised using functional normalisation^89^ in the minfi^88^ package and BMIQ^90^, with 10,000 reference probes for Expectation Maximisation fitting. HPV status was determined using VirusSeq^91^ based on detection of viral gene transcripts.

Beta values for deconvolution associated features, and the signature matrix derived in the previous step, were uploaded to CIBERSORT at https://cibersort.stanford.edu. The data were not quantile normalised due to the potential for global methylation shifts in cancers, and CIBERSORT was run using 1000 permutations. Output files were downloaded as tab-delimited text files and custom parsers were used to import results into R for downstream analysis. FFPE methylation profiles for 42 HNSCC were obtained from Gene Expression Omnibus (Accession GSE38266) using the GEOquery R package, and beta values were BMIQ normalised and analysed using CIBERSORT as described for the TCGA cohort. Wilcoxon’s Rank Sum Tests were used to test for differences in total TIL abundance and TIL subsets.

#### Estimating accuracy of MethylCIBERSORT in tumour deconvolution

In the absence of flow-cytometry based estimates for the different cell types in the analyzed tumours, the estimated fraction of cancer cells from MethylCIBERSORT was compared to sequencing-data based estimates from ABSOLUTE available for 466 HNSCCs from previously published work^29^ using Spearman’s Rank Correlation. Correlations were between ABSOLUTE and other methods of estimating purity*/*immune cell fraction in this subset of tumours; (LUMP, ESTIMATE^76^ and H&E staining assessment of tumour purity (data available in^29^). Residuals were computed by subtracting the method estimate from the ABSOLUTE value. Distributions were compared using Wilcoxon’s Rank Sum Test. Spearman’s Rank Correlation was used to estimate correlations between expression of marker transcripts and MethylCIBERSORT estimates for multiple cellular populations. Where applicable, multiple testing correction was performed using the Benjamini Hochberg approach.

#### Clustering and correlation analyses

Estimates of immune cell fractions in HPV-HNSCC (HPV-transcript negative) were examined for correlations with other infiltrating cell types using Spearman’s Rank Correlation with BH correction for multiple testing. Clustering was carried out using the clusterCons package with 100 iterations using a manhattan distance metric. The most robust number of clusters was then selected.

Differences in the distribution of infiltrating cell types by immune cluster were summarised using mean fold changes and tested using Wilcoxon’s Rank Sum Test with BH-correction for multiple testing.

Differentially expressed genes were identified using Limma-trend and were defined at a threshold of a 2-fold change and BHFDR < 0.01. Pathway analysis was carried out using Ingenuity Pathway Analysis, with findings restricted to experimentally confirmed direct interactions in human cells*/*tissues. Cytolytic activity (CYT) was calculated as the geometric mean of *GZMA* and *PRF1* expression as defined previously^26^. To estimate the contributions of cell population abundances to this, a linear model was fit against log2(CYT) with the different populations as predictors. Wilcoxon’s Rank Sum tests were used to test differences in CYT and CD8:Treg ratios between the immune clusters.

#### Survival Analyses

Multivariate Cox Regression was used to estimate the prognostic utility of clusters derived using infiltration patterns with age and stage as covariates. The survival effect of estimated purity was regressed with the same covariates using a Cox regression with coefficients defined per percent increase in purity.

#### Genomic Correlates

We obtained a list of driver genes inferred by MutSigCV^41^ in TCGA HNSCC cohort from the Broad Institute’s GDAC. GISTIC Copy number estimates thresholded by genes were also obtained from this source. MAF files were obtained from the TCGA data portal. MutSigCV drivers were filtered at a q.value threshold of 0.01 and mutations in this set were tested for differences in frequencies of occurrence using a chi-squared test for differences in proportion. Multiple testing correction was carried out using the Benjamini-Hochberg method. Tables of predicted neoantigens were downloaded from The Cancer Immunome Atlas (http://tcia.at).

#### Benchmarking performance across other tumour types

Signature features were derived from 450k profiles using the aforementioned heuristic (delta-Beta and FDR cutoffs) with a maximum of 100 features per cell type for a wide range of tumour types, using cell lines allocated to the corresponding tissue in GSE6837916^46^ **(Table S10)** and the aforementioned infiltrating cell types **(Table 1)**. These signatures were applied to deconvolve methylation profiles and estimates of purity were derived using TCGA samples for which ABSOLUTE, ESTIMATE and LUMP purity estimates were available^29^.

The cell line data were functionally normalized with the infiltrating cell types described earlier before signature extraction was carried out. 450k data for the aforementioned tumour types were loaded from a pan-cancer freeze derived from SAGE synapse for TCGA pan-cancer (syn2812961) and a custom function was used to extract signature probes and generate methylation percentage matrices for deconvolution with CIBERSORT. CIBERSORT was run as described previously. Correlation and residuals analysis were carried out as described above with MethylCIBERSORT purity estimates vs ABSOLUTE, and between previously published methods and ABSOLUTE. Wilcoxon’s Rank Sum Test with Benjamini-Hochberg correction for multiple testing were used to compare distributions, with these estimates sourced from^29^.

### Pan-cancer analyses of immune cluster assignment

An elastic net model was fit using cellular abundance estimates for HPV-HNSCC using 3 iterations of 5-fold cross-validation to identify the optimal values of lambda and alpha with Kappa values being the selection criterion. The classifier was then applied to MethylCIBERSORT estimates from 18 further tumour types for which corresponding cancer cell line methylation profiles were available^46^ **(Table S10)** to allocate immune cluster. Deconvolution was performed as described above and class allocations were made using the elastic net classifier derived from HNSCC.

For immunoediting analyses, we estimated the number of nonsynonymous mutations encoding at least one immunogenic peptide empirically by summing coefficients across each of six base change contexts as well as the number of non-neoepitope nonsynonymous mutations. Together, these were applied to silent mutation counts in each cancer to derive an expected fraction of neoantigens to nonneoantigens. Comparing the observed fraction to the expected fraction yielded the percentage of neoantigens depleted, and using this in combination with the number of observed neoantigens yielded the count of neoantigen-encoding mutations lost specifically to immunoediting. This was then modelled using a negative binomial framework to estimate the influence of immune cluster on immunoediting.

MAF files for mutations were again downloaded from SAGE synapse for all tumours from the MC3 calling effort (syn7214402). Driver mutations were defined based on pan-cancer MutSig analyses previously published^41^ and binomial GLMs were used to estimate coefficients for mutation frequencies for immune cluster with tumour type as a covariate. Significant genes were defined at BHFDR < 0.1. Survival analyses were performed using data downloaded from Synapse (syn7343873) using Cox proportional hazards regression with stage and cancer type as covariates. Substages were aggregated into stages and only Stages I-IV were considered. Neoantigen abundance and clonality data were downloaded from The Cancer Immunome Atlas^92^.

Negative binomial modelling was used to model all count data, cytolytic activity was modelled using linear models, and binomial GLMs were used to model proportions. Details of covariates, hypotheses and response variables are presented inline. For copy number analyses, SNP6 data were downloaded from the GDC data portal and processed using GISTIC 2.0^93^ on the GenePattern Public Server (arm-level peel off, noise threshold 0.3, FDR < 0.01, driver-gene confidence > 95%) and modelled similarly to mutation data.

### Further resolution of cell-types using expression-based CIBERSORT

RNA-seq data were downloaded from the European Nucleotide Archive for the following datasets: PRJEB11844^94^; GSE60424^95^; and E-MTAB-2319^96^. Kallisto^97^ was used to quantify gene expression with a reference transcriptome consisting of Gencode Grch37 assembly of protein coding and lincRNA transcripts. Data were then modelled using limma trend and the top 50 markers by t-statistics were selected for each cell subset from one versus all comparisons after thresholding with a 2-fold change and FDR < 0.05. These cell types were used to generate a reference profile and CIBERSORT was run to deconvolute samples. For M1*/*M2 macrophage analyses we used LM22 from the CIBERSORT server as the reference. In both cases, Wilcoxon’s Rank Sum Test was used to estimate differences in distributions.

### Analysis of Immunotherapy response

Nanostring data for a panel of immune genes and and exome sequencing data were obtained from Chen et al^54^ and Roh et al^55^ respectively for patients treated using sequential anti-CTLA4 and anti-PD1 checkpoint blockade.

Clustering and machine learning were carried out using the subset of genes intersecting with the Hot-vs-Cold pancancer signature.. 632 bootstrapping was used for hyperparameter tuning and ROC estimation. Negative binomial regression was used to model neoantigen, mutation and subclone numbers, and logistic regression to estimate predictive performance of count data on response.

The number of subclones present in each tumour from the Roh cohort, derived from the EXPANDS algorithm, were obtained from the associated publication^98^. RNAseq data were obtained for aCTLA4 pretreatment biopsies by personal communication with Eliezer Van Allen and genomic data from the associated publication. Data for post-treatment Nivolumab treated melanomas were obtained from _56_.

### Valiation of *EGFR* association with cold tumours

RPPA data were downloaded for TCGA cancers from the TCPA portal. IHC data were derived from^1,7^ for comparison of EGFR protein levels vs TIL levels, previously defined in^1,7^. ssGSEA scores were used to summarise the activity of the glycolytic gene signature (described in) and standard statistical procedures were used to assess interrelationships.

## Acknowledgments

This work was funded by grants from Rosetrees Trust and Debbie Fund (TRF) and a UCL graduate scholarship (AC). We are grateful to Dr Alicia Oshlack for raw DNA methylation data from CD4+ and CD4+*/*FoxP3+ lymphocytes, Elizier Van Allen for aCTLA4 expression data, and Bo Li and Xiaole Shirley Liu for TCR repertoire data. The results published here are, in part, based upon data generated by The Cancer Genome Atlas (TCGA) project established by the National Cancer Institute and National Human Genome Research Institute. Information about TCGA and the investigators and institutions who constitute TCGA research network can be found at http://cancergenome.nih.gov.

## 1 Supplementary Figures

**Supplementary Figure S1:**
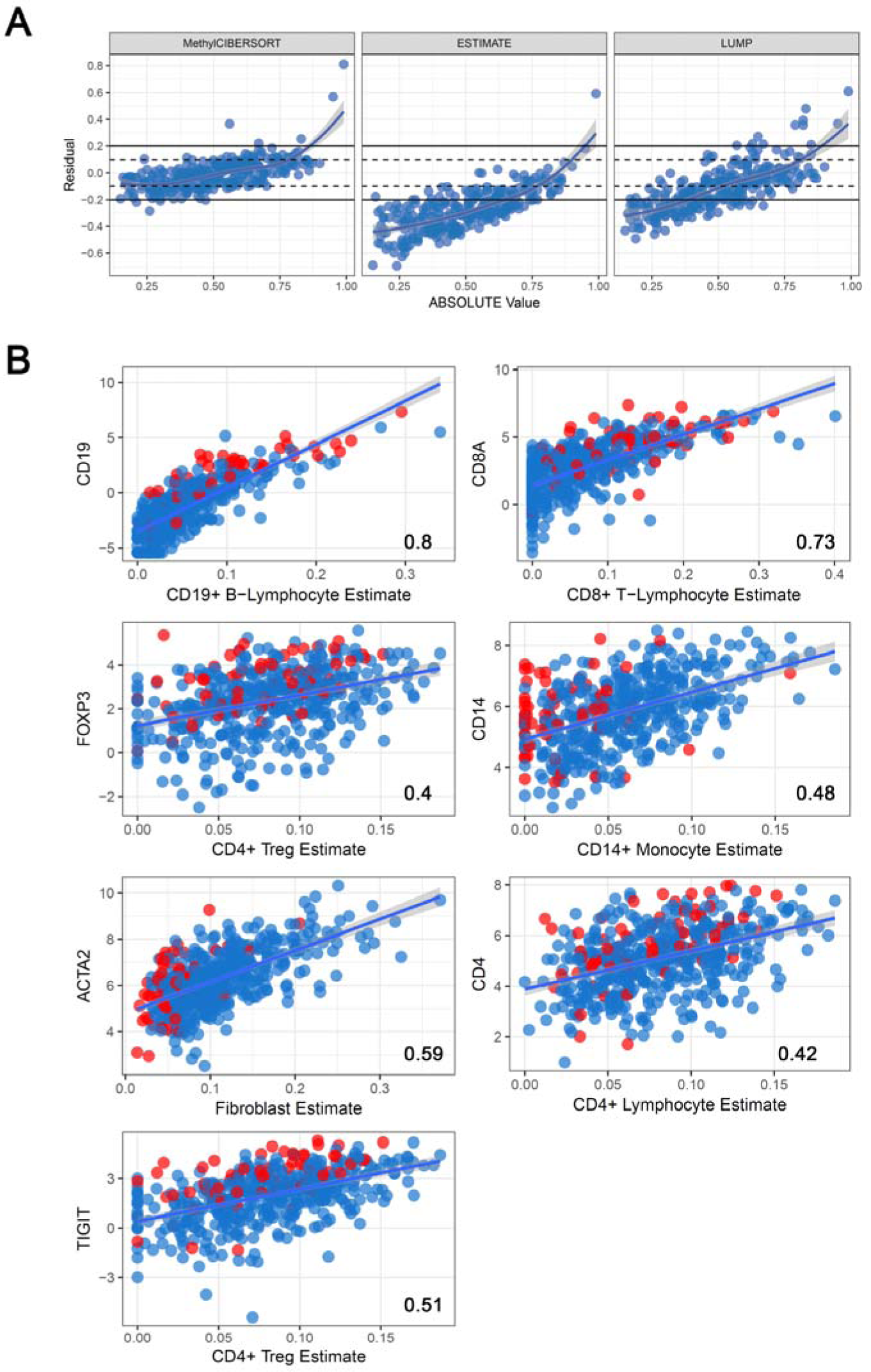
(A) Analysis of ABSOLUTE estimate (x-axis) and error from MethylCIBERSORT, ESTIMATE and LUMP in estimating purity in relation (y-axis). **(B)** Correlations (Spearman’s Rho) between MethylCIBERSORT estimates and marker gene expression in TCGA HNSCC.

**Supplementary Figure S2:**
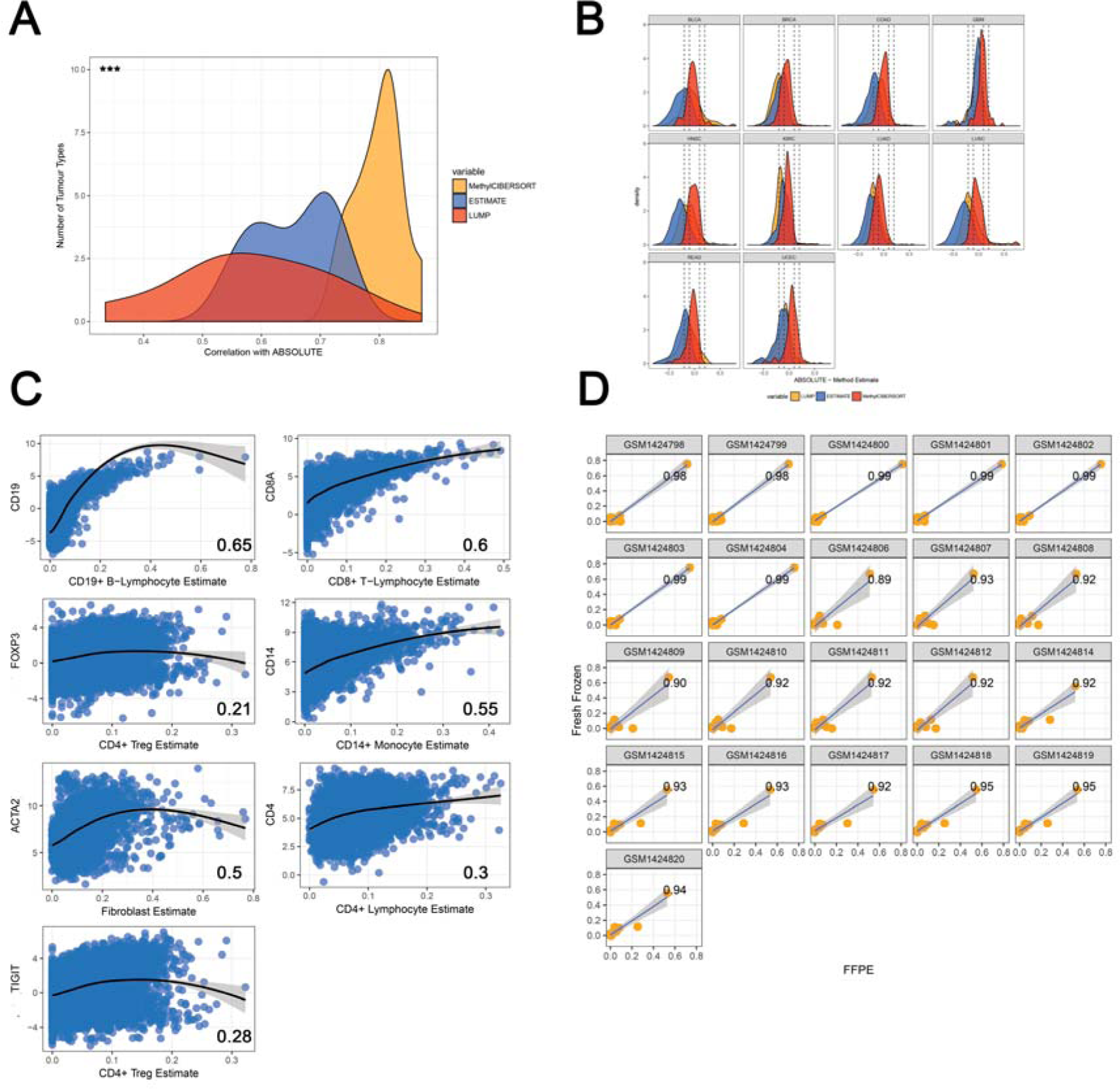
(A) Correlation densities for associations with ABSOLUTE purity for MethylCIBERSORT, ESTIMATE and LUMP across tumour types. **(B)** Density plots showing error relative to ABSOLUTE in individual tumour types for MethylCIBERSORT, ESTIMATE and LUMP. **(C)** Marker correlation plots between MethylCIBERSORT estimates and expression of marker genes. **(D)** Correlation plots for 21 450k methylomes relative from FFPE samples relative to their fresh frozen counterparts.

**Supplementary Figure S3:**
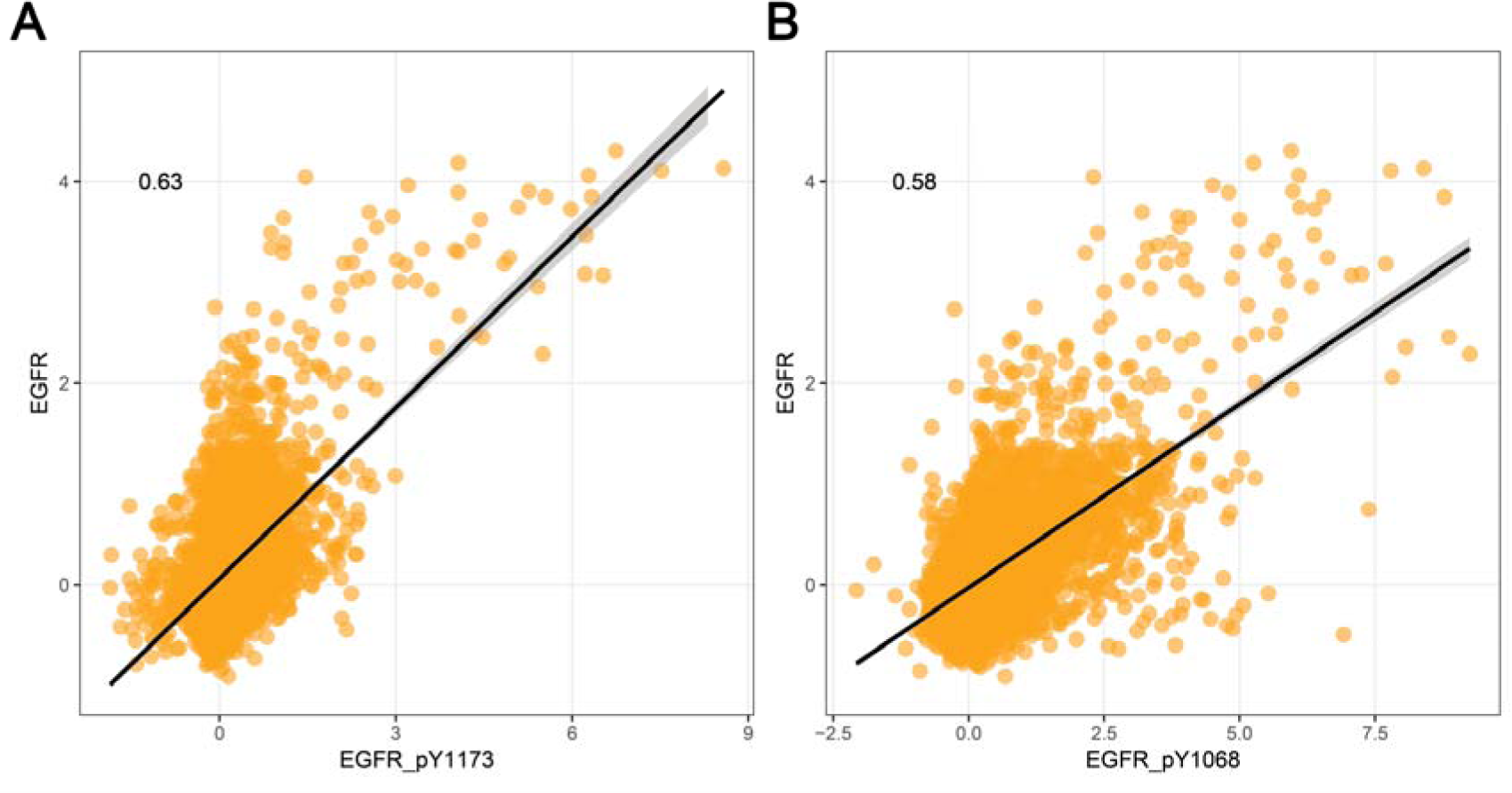
Scatterplots showing association between *EGFR* levels by RPPA and phosphorylation at key activating residues.

## List of Supplementary Tables – Chakravarthy et al

**Table S1** - Genes differentially expressed between HNSCC Hot and Cold clusters.

**Tables S2, S3, S4** – IPA canonical pathway analysis, ontology analysis, and upstream regulator analysis for genes in Table S1, respectively.

**Table S5** - Differentially bound antibodies from RPPA data for HNSCC Hot vs Cold cluster comparison Table S6 – Canonical pathway analysis for pan-cancer hot vs cold transcriptional signature.

**Table S7** – Results of association analysis between immune cluster and mutation frequency.

**Table S8** – Results of association analysis between immune cluster and copy number alteration frequency.

